# Astrocytes locally translate transcripts in their peripheral processes

**DOI:** 10.1101/071795

**Authors:** Kristina Sakers, Allison M. Lake, Rohan Khazanchi, Rebecca Ouwenga, Michael J. Vasek, Adish Dani, Joseph D. Dougherty

**Author notes:** Correspondence to: Joseph Dougherty, PhD Department of Genetics Washington University School of Medicine Campus Box 8232 660 S. Euclid Ave, Saint Louis MO, 63108, USA. (314)362-3774.

## Abstract

Local translation in neuronal processes is key to the alteration of synaptic strength that contributes to long term potentiation and learning and memory. Here, we present evidence that astrocytes have ribosomes in their peripheral and perisynaptic processes, and that *de novo* protein synthesis occurs in the astrocyte periphery. We also developed a new biochemical approach to profile and define a set of candidate transcripts that are locally translated in astrocyte processes, several of which were validated in *vivo* using *in situ* hybridization of sparsely labeled cells. Computational analyses indicate that localized translation is both sequence dependent and enriched for particular biological functions. This includes novel pathways such as fatty acid synthesis as well as pathways consistent with known roles for astrocyte processes, such as GABA and glutamate metabolism. Finally, enriched transcripts also include key glial regulators of synaptic refinement, suggesting that local production of astrocyte proteins may support microscale alterations of adjacent synapses.

**Significance Statement:** Cellular compartments are specialized for particular functions. In astrocytes, the peripheral processes, particularly near synapses, contain proteins specialized for reuptake of neurotransmitters and ions, and have been shown to alter their morphology in response to activity. Regulated transport of a specific subset of nuclear-derived mRNAs to particular compartments is thought to support the specialization of these compartments and allow for local regulation of translation. In neurons, local translation near activated synapses is thought to generate the proteins needed for the synaptic alterations that constitute memory. We demonstrate that astrocytes also have sequence-dependent local translation in their peripheral processes, including transcripts with roles in regulating synapses. This suggests local translation in astrocyte processes may also play a role in modulating synapses.

## Introduction

Astrocytes are known to be indispensable in the development and maintenance of the synapse- a structure described as tripartite due to the critical contribution from peripheral processes of these cells(1). There is evidence that astrocytes determine synapse number(2, 3), and strength(4, 5), and complementary indications that neurons mediate maturation of astrocytes processes(6). Yet, our understanding of these interactions is incomplete.

The localized synthesis of proteins in neurons is instrumental for synaptic modulation(7) and neurite outgrowth(8, 9). Because of the distances neurites must extend to reach a target cell, they have the ability to harbor mRNAs and ribosomes proximal to spines and axons(10, 11) where local translation occurs in a precise spatial, temporal and activity-dependent manner. Additionally, oligodendrocytes also selectively localize some myelin associated mRNAs to specialized domains in their peripheral processes for local translation(12). Outside of the nervous system, localized protein synthesis has been detected in a variety of biological systems, including drosophila oocytes and migrating fibroblasts, suggesting that this phenomenon is a widely used regulatory system(13, 14).

Local translation in both neurons and oligodendrocyte processes suggests that other CNS cells may require this adaptation to compartmentalize functions within the cell. Peripheral astrocyte processes (PAPs) reach lengths comparable to that of some neurites and one astrocyte territory can contact up to 100,000 synapses(15). Further, the dynamic nature of PAPs at synapses requires rapid responsiveness to synaptic changes(5, 16). Therefore, we hypothesized that astrocytes also utilize local translation. We reasoned that several conditions must be met to establish that local translation occurs in astrocytes: First, both ribosomes and mRNA must be present in PAPs. Second, new protein translation must occur in PAPs. Finally, even if protein synthesis is detected, if local translation in astrocytes does indeed support either local adaptation or specialization of function as it does in other cells, then it should be regulated by specific sequence features, and therefore be enriched for specific transcripts. Here, we provide biochemical and imaging evidence for these criteria, and define for the first time the peripheral translatome of the astrocyte.

## Astrocyte ribosomes and mRNA exist in their peripheral processes, *in vivo*

Previous work has indicated that ribosome-like structures exist in PAPs, using electron microscopy(17). We used transgenic mice where ribosomal protein Rpl10a is fused with EGFP (Aldh1L1-EGFP/Rpl10a) specifically within astrocytes to localize the large subunit of ribosomes *in vivo*(18). In cortical astrocytes, we found that EGFP/Rpl10a extends beyond Gfap+ processes and into peripheral processes surrounded by Aqp4, which marks astrocyte membranes and vascular endfeet(19) (Fig. 1A,B). As translation requires both large and small subunit, we confirmed peripheral localization of endogenous small ribosomal subunits via immunofluorescence (IF) against Rps16 (Fig. S1), indicating the tagged Rpl10a is not simply mislocalized in the Aldh1L1-EGFP/Rpl10a mice. We also asked if we could detect tagged ribosomes in perisynaptic PAPs using a recently-developed approach leveraging stochastic optical reconstruction microscopy (STORM), to allow for ultrastructural level resolution compatible with multicolor fluorescent labeling.(20) Astrocyte EGFP/Rpl10a was found within 100 nm of synapses, which are defined by apposition of pre-and post-synaptic markers (Fig. 1C). This confirms the earlier EM work indicating ribosome-like structures are found in the peripheral processes of astrocytes, and suggests that the EGFP tag might be used for purification of ribosome bound transcripts, if mRNA is indeed present and translating on these ribosomes.

**Figure 1:**
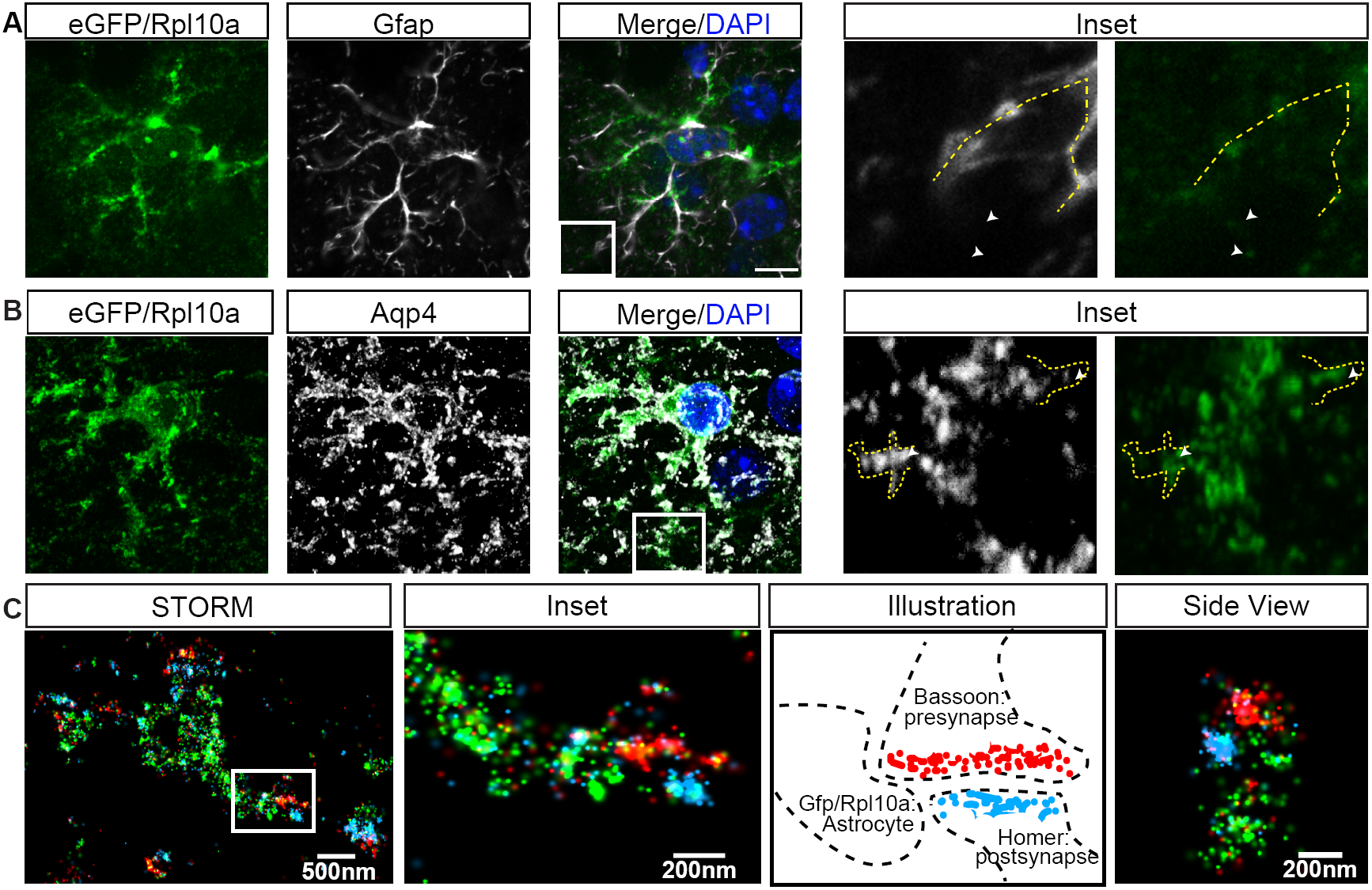
**Subcellular localization in cortical astrocytes shows ribosomal proteins in peripheral processes and in close proximity to synapses** (A-B) Confocal IF of a cortical astrocyte, stained with Gfap(A) or Aqp4(B)(white). EGFP/Rpl10a(green) shows ribosomal tag extending throughout astrocyte, past Gfap+ processes (yellow dashed line) and within fine processes highlighted by Aqp4. Scale bar=10μm. (C) STORM imaging showing an EGFP/Rpl10a (green) filled astrocyte process proximal to synapses(as illustrated, these are defined by apposition of Bassoon(red), and Homer(blue)). Inset of box on left, and side view is a 90 rotation of a second synapses, again showing EGFP/Rpl10a puncta surrounding a synapse.

Therefore, we next sought to confirm that at least some mRNAs are found in PAPs. We chose to focus on the localization of *Slc1a2* (Glt-1) for two reasons: first, Glt-1 protein is highly expressed in PAPs(21) and second, previous colormetric *in situ* hybridization (ISH) suggested its peripheral localization(22). As both Aqp4 antibody and the pan astrocyte Aldh1L1-EGFP/Rpl10a were suboptimal for easily tracking the processes of single astrocytes, we developed a viral method to sparsely GFP-label astrocytes in non-transgenic mice (Fig. 2A). Using this approach, we confirmed that *Slc1a2* mRNA is found throughout the astrocyte, including in distal GFAP negative PAPs (Fig. 2B).

**Figure 2:**
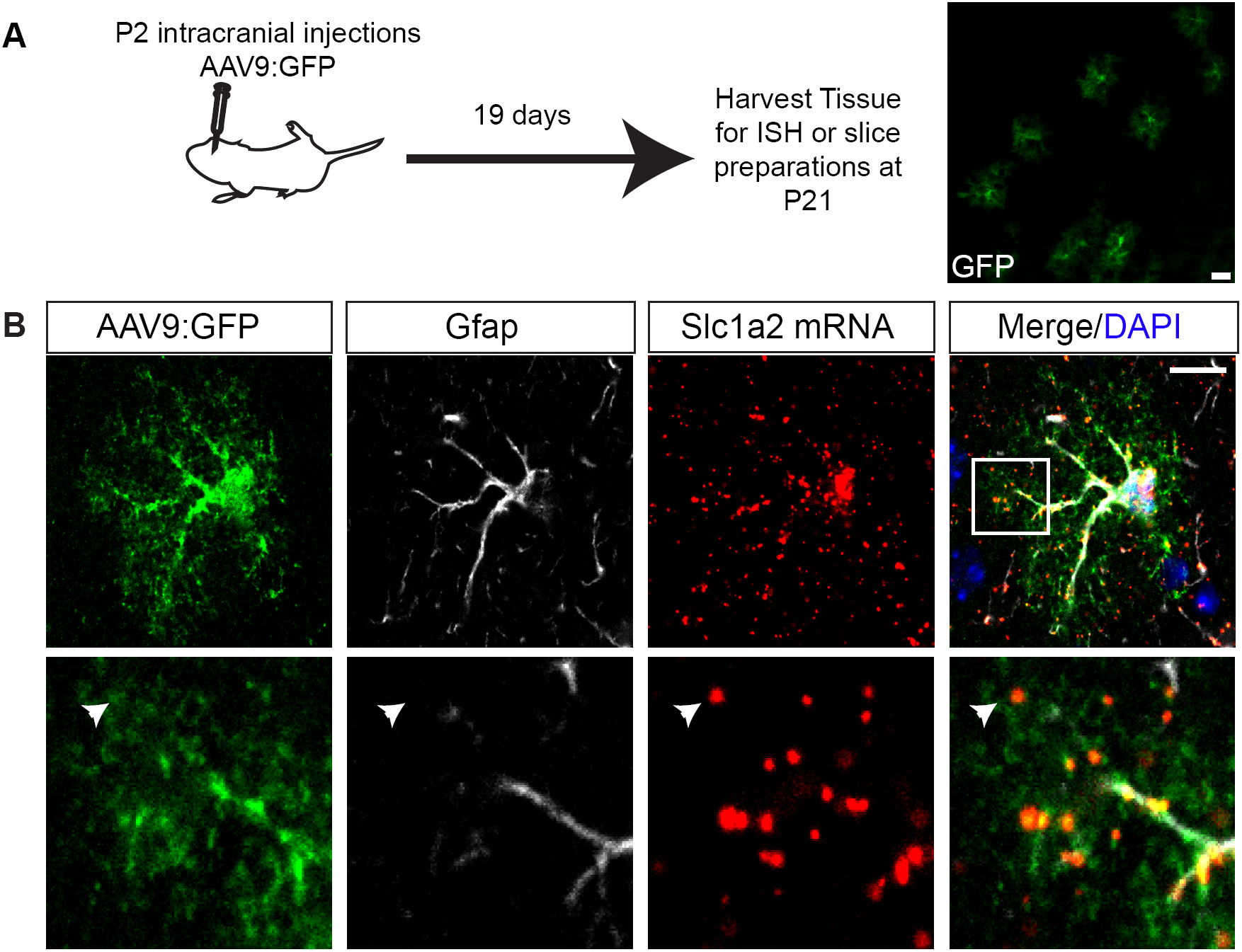
**Method for sparse astrocyte labeling and FISH of *Slc1a2* reveals its peripheral localization.** (A) Early postnatal pups were injected transcranially with AAV9 expressing GFP, resulting in sparse labeling of cortical astrocytes after 3 weeks. Scale bar=20μm. (B) Confocal IF of a GFP-labeled astrocyte shows extension of *Slc1a2(red*) FISH into fine, Gfap(white) negative peripheral processes, scale bar=10μm.

## Astrocyte processes synthesize proteins in their distal tips

After demonstrating the presence of both ribosomes and mRNA in PAPs, we next tested whether distal translation occurs by using puromycin to label actively-translated peptides in acute brain slices (Fig. 3A). Puromycin, a tRNA structural analog, incorporates into translating ribosomes and puromycylates the growing peptide(23). Using an antibody against puromycin for IF in sparsely labeled astrocytes (Fig. 2A, S2A), we were able to detect nascent translation within PAPs, which was blocked by pretreatment with another translation inhibitor, anisomycin, indicating that labeling requires active translation (Fig. 3B). We quantified the total translation occurring throughout the astrocyte by measuring the fluorescence intensity of puromycin in the astrocyte (determined by colocalization with GFP) at increasing radii from the nucleus of the astrocyte. We empirically determined that a 27um radius captures the majority of the astrocyte, without substantially extending to another astrocyte territory (Fig S2B), which is consistent with previous data regarding a mean astrocyte diameter of 56 microns(24). Given the short duration of incubation, we concluded that puromyclated peptides in PAPs were made locally rather than transported. Quantification of puromyclation in astrocytes, indicates that on average 73% of translation in an astrocyte occurs > 9 microns from the nucleus (Fig. 3C), and does not taper significantly as it extends to the periphery.

**Figure 3:**
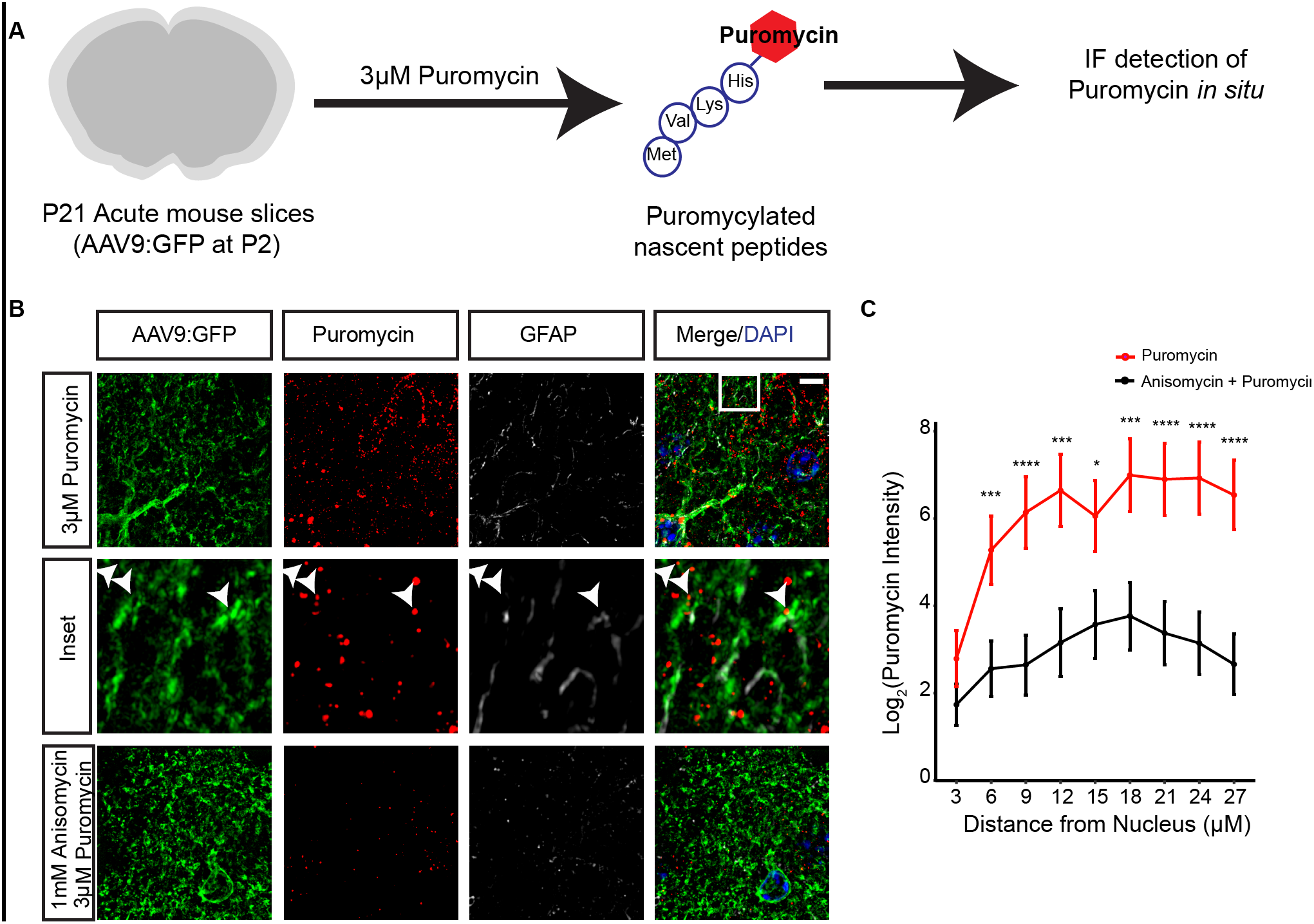
**Peripheral ribosomes are actively translating in astrocytes** (A) Acute slices(300 μΗ) from P21 mice with sparse astrocyte labeling(generated as in Fig. 2), are treated for 10 minutes of puromycin, with or without 30 minutes of anisomycin pretreatment, then fixed, sectioned, and processed for IF against puromycin(red) and GFP(green). (B) Max projection super-resolution Structured Illumination Microscopy(SIM) to detect puromyclation(red) of synthesizing proteins in a GFP-labeled astrocyte shows translation occurs in peripheral processes, and is blocked by pretreatment with anisomycin. Scale bar=10μm. (C) Quantification of puromycin intensity (only puromycin pixels that were in astrocytes labeled with GFP were measured) at increasing radii from the nucleus indicates robust translation occurs in PAPs. Repeated measures ANOVA revealed main effects of condition F(2, 138)=9.694, p=0.0001 and distance F(8, 1104) = 19.023, p < 2E-16 with a significant interaction between condition and distance F(16, 1104) = 2.019, p=0.01. Data represented as mean +/-SEM. Asterisks represent one-sided T-tests, *post hoc.* **** p < 0.001, ***p < 0.005, **p < 0.01. N(cells) = 54(puromycin), 48(anisomycin+puromycin).

## PAP-TRAP reveals hundreds of enriched ribosome-bound transcripts in PAPs

In neurons, local translation appears to be transcript-specific, often generating proteins with synaptic roles. If translation in PAPs does indeed have a physiological role, then a specific subset of astrocyte transcripts should be translated there. In other words, the localization of RNAs to the periphery should be sequence dependent, and consistent with the known roles of the PAP in glutamate homeostasis and other processes. To capture ribosome-bound mRNA from PAPs, we applied translating ribosome affinity purification(TRAP) to a synaptoneurosome(SN) preparation from Aldh1L1:EGFP/Rpl10a mice, a method we have abbreviated ‘PAP-TRAP’ (Fig. 4A). While SN fractions have been previously studied largely for containing synaptic structures, previous work has indicated at least one peripherally localized astrocyte mRNA is present in them(25) and our preliminary RNA sequencing of this fraction suggested substantial contribution of mRNA from non-neuronal *cells(data not shown*). Here, immunoblot confirmed depletion of nuclear protein LaminB2 and enrichment, as expected, of Psd95 in the SN fraction (Fig. 4B). Additionally, we show the fraction contains Ezrin, a constituent of PAPs(26), and astrocytic EGFP/Rpl10a (Fig. 4B). Thus, we used this affinity tag to harvest astrocyte ribosomes from the fraction, and performed RNA-seq on the PAP-TRAP sample and three comparison samples (Table S1) to identify PAP-enriched transcripts (Fig. 4A). Our analysis identified 224 transcripts that are significantly enriched on PAP ribosomes (Fig. 4C, Table S2), including *Slc1a2*, and 116 that are depleted(‘Soma’ transcripts, Fig. 4C, Table S3). These data establish that PAP-TRAP samples contain a distinct profile of ribosome-bound mRNAs, which includes the significant enrichment of mRNAs for proteins, such as Slc1a2(Glt-1), known to be found in PAPs (Fig. 4D). Unfortunately, PAPs have no sufficiently distinctive stable structure equivalent to a post-synaptic density that can be used to readily verify their presence in SN fractions by electron microscopy. Therefore, we instead focused on using an independent anatomical method *in vivo* to validate the novel predications of the PAP-TRAP purification experiment. Using FISH, we successfully confirmed the peripheral localization of several of these new candidates (*Cpe, Clu, Glul*, and *Sparc*) (Fig. 5).

**Figure 4:**
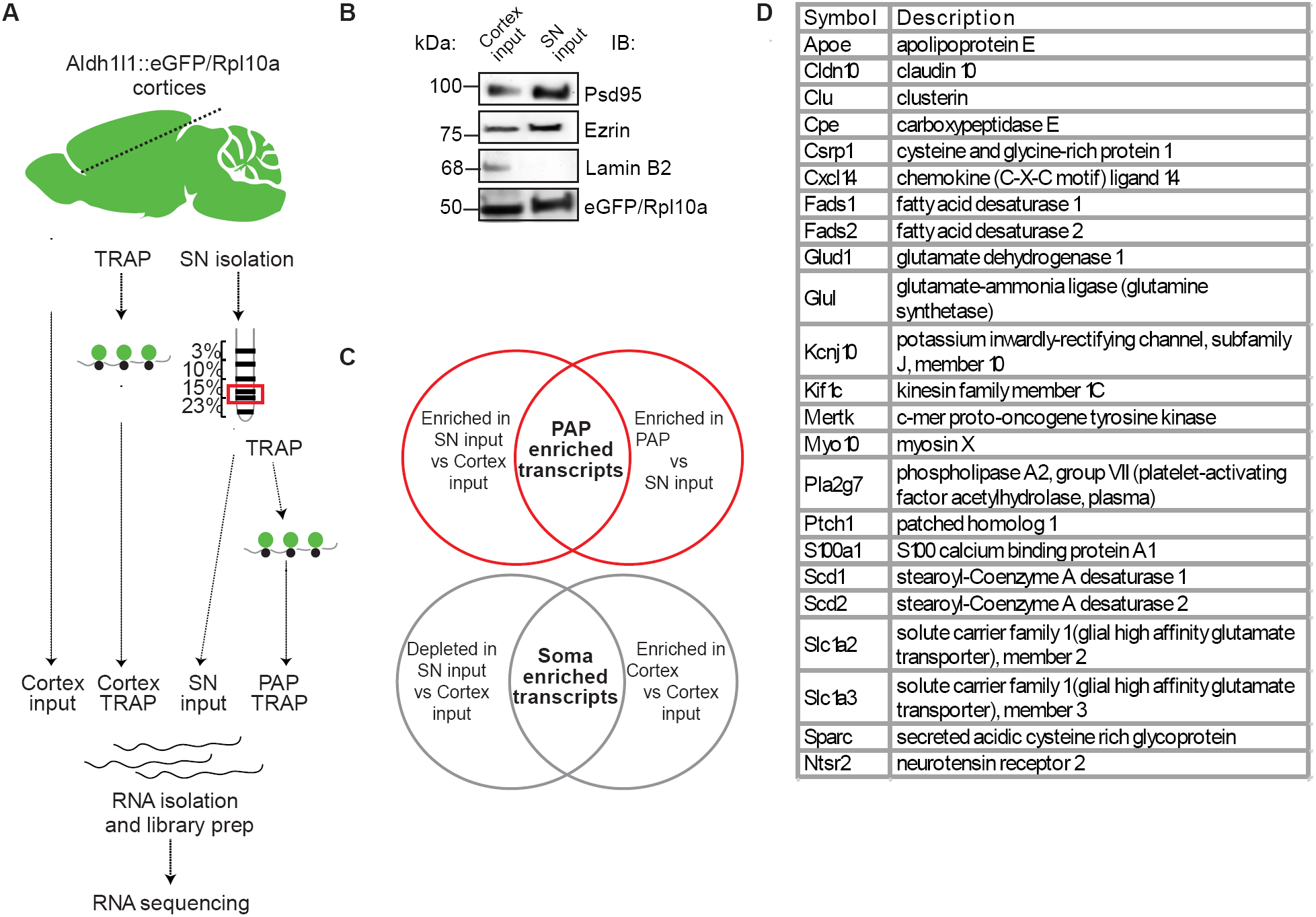
**Identification of peripherially enriched transcripts** (A) Diagram of experimental steps in PAP-TRAP and comparison samples for RNA-seq (B) Representative immunoblots for input and SN fraction confirms enrichment of synaptic and PAP proteins, depletion of nuclear proteins (LaminB2) and presence of EGFP/Rpl10a (C) Diagram of analytical strategy for defining PAP and Soma enriched transcripts (D) Examples of PAP-enriched transcripts, including those related to Glutamate metabolism(Slc1a2, *Slc1a3, Glul*), fatty acid *synthesis(Fads, Scd*), and interesting signaling molecules(Ptch1, *Sparc, Ntsr2*).

**Figure 5:**
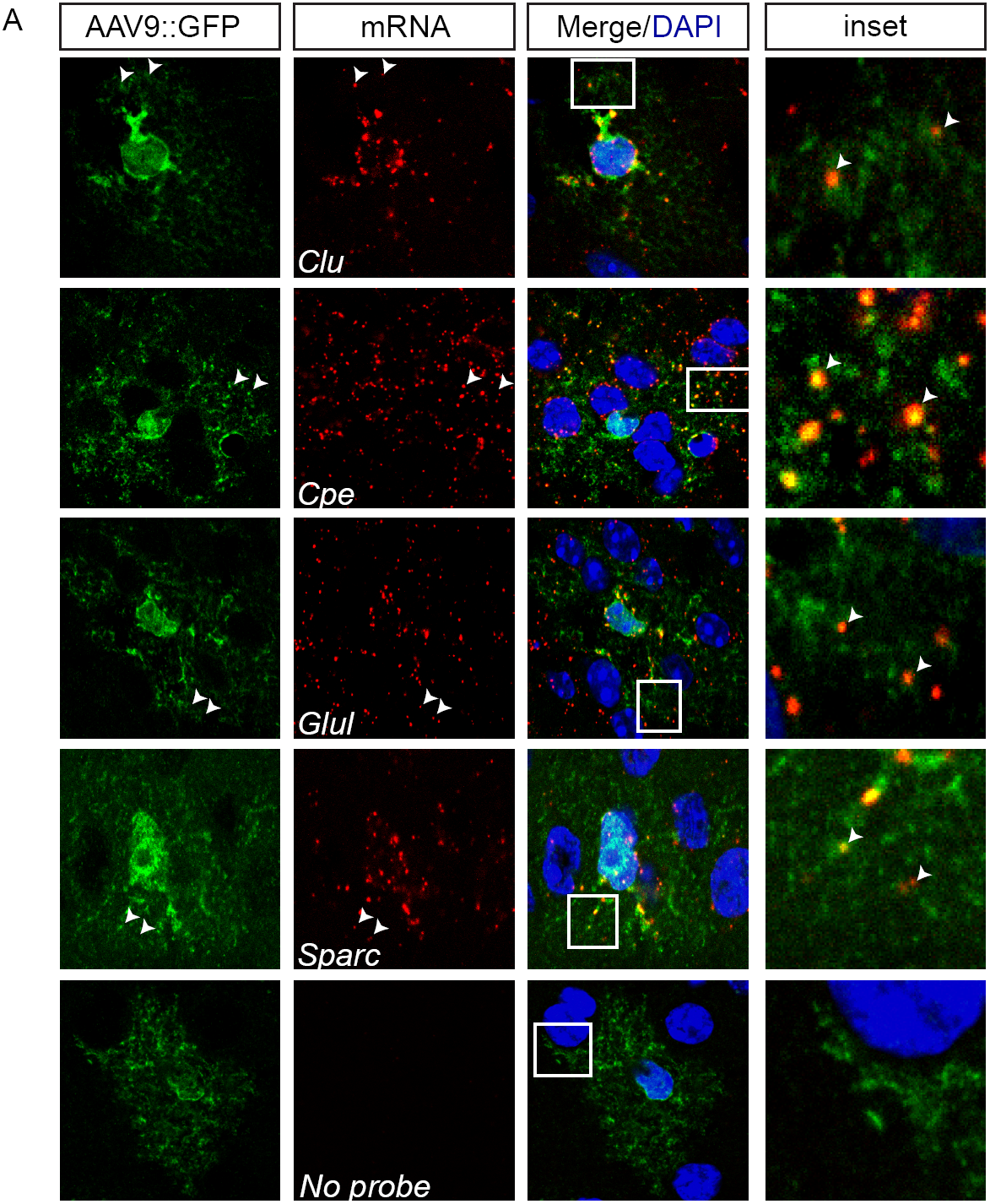
***In vivo* validation of PAP-TRAP candidates** (A) FISH on GFP-labeled astrocytes confirms presence of PAP-TRAP identified mRNAs(Cpe, Clu, Glul and Sparc) in peripheral processes. Cortical astrocytes were labeled as described in Fig 2. Images are representative of three independent experiments with each probe.

## PAP-enriched transcripts suggest physiological roles for local translation including modulation of neurotransmitter metabolism

To understand what purpose local translation might serve in astrocytes, we did a pathway analysis to identify consistent biological functions of the PAP enriched transcripts (Fig. 6A). We found that PAP-localized transcripts had overrepresentation of genes mediating Glutamate and GABA metabolism, consistent with PAP functions of Glutamate transport(Slc1a2, *Slc1a3*)(21, 27) and metabolism(*Glul*). We also identified a set of enzymes representing multiple steps in a pathway for biosynthesis of unsaturated fatty acids (*Scd1, Scd2, Fads1, Fads2, Elovl5, Hadha*) suggesting there may be some local regulation of fatty acid production either for signaling or membrane use. We also identified several motor and cytoskeletal proteins(e.g *Kif1c, Myo1D*) local translation of which could play a role in morphological remodeling of astrocyte processes. Finally, we also noticed an enrichment for genes and members of gene families known to regulate synapse number(Mertk, *Sparc, Thbs4*)(28–30) suggesting that synapse formation and elimination may be mediated in part by local translation of signaling molecules in astrocytes. Importantly, astrocyte soma transcripts have largely non-overlapping GO categories with PAP transcripts and are enriched for transcription regulators and amino acid catabolism (Fig. S4).

**Figure 6:**
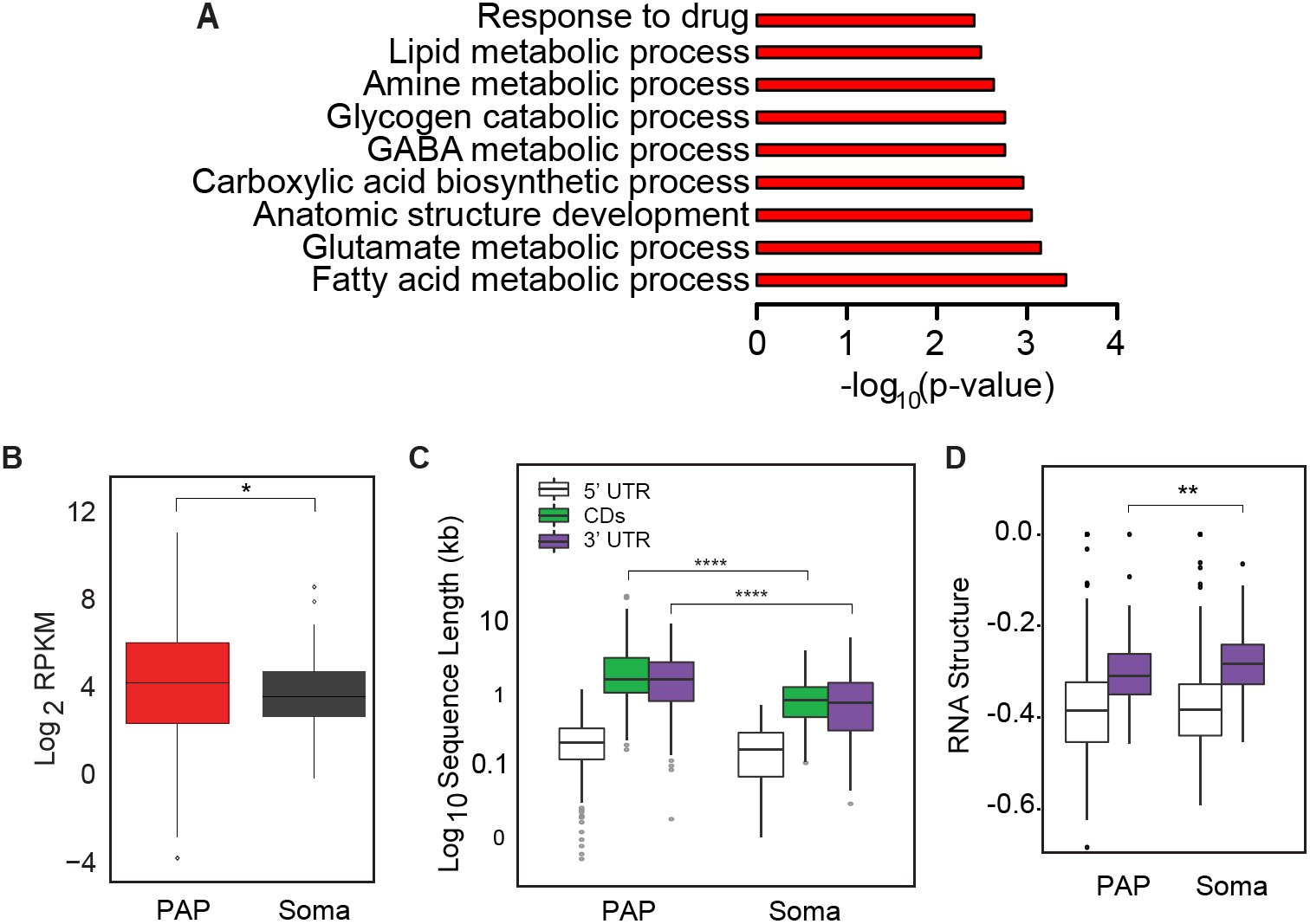
**Pathway and sequence analysis on PAP-enriched transcripts** (A) Representative significant GO terms for PAP-enriched transcripts, hypergeometric test with Benjamini-Hochberg correction(B-H) (B) Quantification of expression of PAP-enriched and Soma-enriched transcripts indicates Soma transcripts have lower median expression in cortical astrocytes(Wilcoxon test, B-H corrected, *p<0.05) (C) Quantification of length of PAP and Soma transcripts indicates PAP-enriched transcripts have longer 3’UTRs(Wilcoxon test, B-H corrected, ****p<0.0001). (D) RNA structure-score (minimum free energy of most stable predicted structure, normalized to length), indicates PAP-enriched transcripts have more stable 3’UTR secondary structures (Wilcoxon test, B-H corrected, **p<0.01, lower values are more stable).

## PAP-enriched transcripts have longer 3’UTRs and are higher expressed than Soma-localized transcripts

For dendritically localized transcripts, as well as the localization of β-actin in fibroblasts, sequence-specific features are commonly found within the 3’UTR that mediate their localization(31–33). Therefore, we asked whether PAP-enriched transcripts contain sequence-specific features or characteristics that may mediate localized translation. We found no differences in individual nucleotide or GC content (Fig. S5). However, PAP-enriched transcripts were significantly higher expressed in cortical astrocytes compared to ‘Soma’ transcripts (Fig. 6B). In neurons, locally translated mRNAs, often sequestered by RNA-binding proteins(RBPs) into granules for transport and protection, have significantly longer 3’UTRs(34). This may enable more sequence motifs or secondary structure for binding RBPs. We found that PAP-enriched transcripts are also significantly longer, specifically in their 3’UTRs (Fig. 6C). Additionally, RBPs may recognize secondary structure rather than specific sequence. We used the RNA-folding algorithm, ViennaRNA(35), to investigate the stability of transcript 3’UTRs in the PAP and Soma transcripts. Interestingly, we found that PAP-enriched 3’UTRs are predicted to have more stable secondary structures (Fig 6D). The greater length and structure suggests that sequences in 3’ UTRs may also serve to regulate the localized enrichment of specific mRNAs in PAPs.

## Discussion

Here we provide, to our knowledge, the first evidence of localized translation of specific transcripts in astrocytes. We utilize a novel biochemical method to enrich for ribosome-bound transcripts in PAPs and provide independent *in vivo* validation of several of the transcripts, confirming the PAP-TRAP method is capable of selecting for peripherally enriched astrocyte mRNA. Additionally, we have provided evidence that translation indeed occurs outside of the astrocyte soma, as indicated by labeling of new protein synthesis with a brief pulse with puromycin. With this approach we have confirmed that peripherally localized puromycylated peptides extend to the very distal processes of the astrocyte.

Our data also indicate that PAP-enriched candidates have a median higher expression and a significantly longer and more stable 3’UTR than those transcripts not enriched in the PAP. These findings suggest that localization information exists in the 3’UTR, as is the case with many dendritically-localized transcripts(32). Understanding of the mechanisms governing this localization will provide insight to the phenomenon and opportunities to test the physiological relevance of localized translation in astrocytes. Interestingly, we also detected a few neuron-derived transcripts in the PAP-localized transcripts(Table S2), but we conclude that the removal of these transcripts from the data does not change the overall conclusions of the PAP-localized transcripts (Fig. S7, Table S4). The possible mechanisms for this signal may reveal interesting biology and are of interest for future study.

We also note that our current list of locally translated candidates includes several disease genes, most notably the Alzheimer’s associated gene *Apoe* appears to be robustly enriched in the PAP-TRAP sample. Further, at least two transcripts reported to be disrupted using TRAP in astrocytes in Amyotrophic Lateral Sclerosis (ALS) models (*Heyl*, and *Vim*)(36), as well as *Slc1a2,which* is downregulated in the motor cortex of ALS patients(37), are enriched in the PAP-TRAP sample. It is possible that dysregulation of local translation may contribute to disease progression and vulnerability of adjacent neurons. The PAP-TRAP method should be readily adaptable study this possibility in mouse models of neurological disorders.

While local translation in neurons has been described for decades, many years later very few studies have been able to address whether or not local translation specifically, rather than local translation more generally had a physiological role. Local translation was first shown to be important in the induction of long term potentiation (LTP), whereby physical separation of the dendritic layer in the hippocampus was sufficient to induce LTP, but translation was required as LTP was blocked by pretreatment with a protein synthesis inhibitor (7). Moreover, mutation of the dendrite-targeting element of *CamKIIα* resulted in decreased performed in spatial memory tasks such as the Morris Water Maze(38). Though we do not yet have the tools to directly probe the physiological role of astrocyte translation, our pathway analyses provide interesting clues. We found that the transcripts for secreted protein precursors are found in PAPs (*Clu, Cpe* and *Sparc*), all of which have been identified as enriched in astrocyte-conditioned media by mass spectrometry(39). This suggests that these glial derived soluble signaling molecules may be locally synthesized before their extracellular release. Further, we noted transcripts for multiple proteins with known roles in maintaining synapse numbers were enriched on PAP ribosomes (*Sparc, Mertk, Thbs4*). It has been shown that Sparc negatively regulates excitatory synaptogenesis by antagonizing Hevin(29). Further, Mertk is a required engulfment receptor for astrocyte-mediated synapse elimination, *in vitro* and *in vivo*(28). Together, these findings provide evidence consistent with the exciting possibility that localized translation might have a role in a localized astrocyte-mediated synaptic modulation. Overall our data overwhelmingly indicate that astrocytes are capable of a sequence-regulated localized translation. We posit here that, whereas neurons are thought to require localized translation to compensate for the great length of their dendrites and axons, astrocytes contacting multiple synapses may use local translation to allow for them to respond to or modulate the activity of specific synapses.

## Materials and Methods

### Animals

All procedures were performed in accordance with the Washington University Institutional Animal Care and Use Committee (IACUC) and the Animal Studies Committee (ASC). Mice were maintained in standard housing conditions with food and water provided ad libitum. B6.FVB-Tg(Aldh1L1-EGfp/Rpl10a)^JD130Htz^/J animals(40) were bred to pure C57BL/6J mice (Jackson strain # 000664) to obtain litters of GFP+ and GFP-animals.

For sparse labeling of astrocytes for microscopy, AAV9-CBA-IRES-GFP injections were performed in P2 pups. 2μl of 10^12^ vector genome (vg) / mL virus was injected bilaterally in the cortex, 1.5mm from the midline in two regions: 1mm caudal to bregma and 2mm rostral to lambda. Injections were performed with a 33g needle (Hamilton #7803-05) with a 50μl Hamilton syringe (#7655-01). Mice were aged until 3-4 postnatal weeks prior to CO2 euthanasia and transcardial perfusion with PBS, followed by 4% paraformaldehyde in PBS.

### Acute Slice Preparation and Drug Treatment

Acute cortical slices (300μm) were prepared in artificial cerebrospinal fluid (aCSF, in mM: 125 NaCl, 25 glucose, 25 NaHCO3, 2.5 KCl, 1.25 NaH2PO4, equilibrated with 95% oxygen-5% CO2 plus 0.5 CaCl2, 3 MgCh; 320 mosmol). 3μM Puromycin (Tocris #40-895-0) was added to slices and allowed to incubate for 10 minutes at 37C. 1mM Anisomycin (Sigma #A9789) in aCSF was added to slices 30 minutes before Puromycin treatment at 37°C.

### Immunofluorescence (IF)

40μm sections were cryosectioned after 4% paraformaldehyde transcardial perfusions. Sections were incubated with either: Rabbit GFAP (Dako, 1:1000), Rabbit Rps16 (Sigma, 1:1000), Rabbit Aqp4 (Santa Cruz, 1:100), Mouse Puromycin (Kerafast, 1:1000) followed by detection with appropriate Alexa conjugated secondary antibodies (Invitrogen) and nuclei were counterstained with DAPI. For dual FISH/IF, GFP was detected with Chicken GFP (Aves) at 1:1000, and detected as above. For STORM IF: 14μm sections were cryosectioned after flash freezing on dry ice and images as previously described(41). Cryosections were immunolabelled with primary antibodies, Chicken anti-GFP (Invitrogen), Mouse anti-Bassoon (SAP7F407, Novus Biologicals) and Rabbit anti-Homer1 (Synaptic Systems, Germany) followed by anti-chicken, anti-rabbit and anti-mouse secondary antibodies raised in donkey (Jackson Immunoresearch). Secondary reagents were custom conjugated with acceptor and reporter fluorophore dye pairs Cy3-Alexa647, Cy2-Alexa647 and Alexa405-Alexa647. Immunolabelled sections were overlaid with a buffer containing 100mM Tris-HCL pH8.0, 150mM NaCl and containing an oxygen scavenging system comprised of glucose, glucose oxidase, catalase and the reducing agent 2-mercaptoethylamine (Bates *et al.*, 2007). After removing excess buffer, edges of the coverslip were sealed with nail polish before imaging. Representative images are from at least triplicate experiments.

### Fluorescent In Situ Hybridization (FISH)

FISH was performed on 14μm slide-mounted cryosections after 4% paraformaldehyde fixation of either Aldh1l1:eGFP/Rpl10a or AAV9-CBA-IRES-GFP animals. Digoxigenin-labeled antisense probes were made from purified PCR products containing a T7 promoter site in the appropriate orientation. In vitro transcription was performed using T7 polymerase (Promega #P2075) and Dig RNA labeling Mix (Roche #11277073910) according to manufacturer’s instructions. Primers for probe template were as follows:

*Slc1a2:* 5’ GTTCCCTCCCAATCTGGTAGA 3’, 5’ *TAATACGACTCACTATAGGGCCGTGGCTGTGATGCTTATT* 3’

*Cpe:* 5’ TTTTCCAAAGCTTGGCTCGC 3’, 5’ TAATACGACTCACTATAGGGTGTGATTGCCAGGTAGCCAG 3’

*Clu: 5’* GCTCTGGAGGACACTAGGGA 3’, 5’ TAAT ACGACTCACTATAGGGCCCAT AGT GGGAGGGAGACA 3’

*Sparc*: 5’ GCCTGGATTCCGAGCTG 3’, 5’ TAATACGACTCACTATAGGGCCTCCAGGGCAATGTACTTGT 3’

*Glul:* 5’ CGGGCCTGCTTGTATGCTG 3’, 5’ TAATACGACTCACTATAGGGTCTTCTCCTGGCCGACAGTCC 3’

Hybridization was performed at 65C overnight in a humid chamber, followed by washes and H_2_O_2_ block as described in(42).

Detection was performed using Sheep Anti-Dig-POD (Roche #11207733910) followed by Tyramide Signal Amplification Cyanine 3 Tyramide (PerkinElmer #NEL704A001KT) as per manufacturer’s instructions. Slides were then incubated with antibodies against additional specific proteins (see IF above) for dual FISH/IF.

### Microscopy

Confocal microscopy was performed on an UltraVIEW VoX spinning disk microscope (PerkinElmer) or an AxioImager Z2 (Zeiss). SIM microscopy was performed on a Nikon n-SIM microscope.

STORM images were acquired on a custom built single molecule-imaging rig. Briefly, the imaging rig was constructed on an active dampening vibration isolation table (Newport) and around a Nikon TiE inverted microscope. Lasers, 642nm (Vortran laser, CA), 561nm, 488nm (Sapphire, Coherent Inc., CA) and 405nm (Cube, Coherent Inc., CA) were shuttered individually via mechanical shutters as well as an acousto optical tunable filter (AOTF, Crystal technologies, PA). Lasers were combined, expanded, collimated and focused at the back focal plane of the microscope. The diameter of the laser beam was cut using a field diaphragm to just cover a 256x256 pixel area of the CCD camera. A mechanical translation stage enabled objective type total internal reflection (TIR) illumination via a 100X, 1.4NA objective (UPlanSApo, Olympus). The Nikon perfect focus system was used to lock focus at the coverslip surface during imaging. A quad band dichroic mirror zt/zet-405/488/561/640rpc (Chroma Technology, VT) was used to excite with laser light and fluorescence collected by the same objective was filtered using et705/72m emission filter for Alexa647. Images were acquired on a back illuminated EM-CCD camera (DU-897, iXon+, Andor Technology, UK), using custom software. 3D astigmatism imaging was performed as described before by placing a cylindrical lens in front of the camera. Sparse single molecule images were acquired at 60Hz frequency using an imaging and activator laser sequence: one frame of weak activator laser (561/488/405) followed by three frames of 642nm laser at 560W/cm^2^, repeated as a train for each activator dye-antibody combination. The amount of activation lasers was adjusted to ensure sparse single molecule events in each camera frame. Raw image stacks were analyzed to determine the centroid positions of fluorescent intensity peaks and these STORM localizations were rendered as images using custom software.

### Quantification of imaging data

To determine the distribution of translation occurring in the astrocyte, we drew concentric rings around the nucleus with increasing radii of 3μm. We empirically determined that a circle with a radius of 27μm encompasses the entire cell, without approaching another astrocyte’s territory. The results of this quantification is demonstrated in Fig 3. Analysis was conducted in ImageJ and described in Figure S2. Experiments were conducted in triplicate. Cells counted are reported in the legend for figure 3.

### Synaptoneurosome (SN) preparation

SN fractions were isolated using a modified protocol(43). Three cortices (including hippocampi) were pooled from Aldh1l1:eGFP/Rpl10a mice and homogenized in 3.5mL of: 5mM Tris pH 7.5 and 250mM sucrose supplemented with 0.5mM DTT, 100μg/mL cycloheximide, 1mM tetrodotoxin (Tocris #1069), 40units/mL rRNAsin (Promega #N2511), 20units/mL SUPERaseIn (Ambion #AM2694) and 1 tablet/10mL cOmplete Mini EDTA-free protease inhibitor cocktail (Roche #4693159001). Homogenization was performed in a glass dounce homogenizer (7mL max volume) with 10 strokes each of the loose and tight pestles. The homogenate was first spun at 1000xg in a Sorvall RT7 plus centrifuge for 10 minutes at 4°C without brake. 2mL of the 2.5mL supernatant was layered onto a sucrose percoll gradient as described (43) in Seton open-top tubes (#5042) and spun at 32,500xg for 5minutes in a Sorvall RC-5C without brake. The SN band was collected by puncturing the bottom of the tube, allowing the bottom milliliter to drip out and collecting the next 3.5mL, which are the combined SN fractions as described(43). 200μ1 of the SN fraction was used as the SN input sample. The remaining homogenate supernatant (~500μl) was spun at 20,000xg for 15mins at 4C after incubation with 1% NP-40(Sigma) and 30mM DHPC (Avanti). 60ul of the supernatant was added to 140μl of 0.15M KCl supplemented as the homogenization buffer, and was set aside for use as the Cortex input sample. The remaining supernatant and the SN fraction were used for either TRAP or Western blotting.

### Translating ribosome affinity purification (TRAP)

TRAP was performed on 3 replicates of SN fraction (PAP TRAP sample) and cortex homogenate (cortex TRAP sample), modified from protocols as described(40, 44). Briefly, biotinylated-Protein L coated streptavidin magnetic beads were coupled with 2 monoclonal mouse Anti-EGFP antibodies(40). Samples were added to conjugated beads and incubated on a inverted mixer for 3 hours at 4°C. Beads were then washed 3 times with a high-salt buffer (10 mM HEPES at pH 7.4, 350 mM KCl, 5 mM MgCh, 1% NP-40, 0.5 mM dithiothreitol, 100 μg/mL cycloheximide). RNA purification was performed on eluted beads and respective input samples in parallel using Qiagen RNEasy MinElute kit. RNA quality and concentration was assessed using PicoChips on the Agilent BioAnalyzer. As large subunit was tagged, representative traces (Fig S6) containing both 18S(small subunit) and 28S(large subunit) ribosomal RNA confirm that 80S ribosomes were harvested from the SN fraction and comparison groups. All samples used in this experiment had RIN scores > 8 for RNAseq library preparation.

### Library preparation

Double stranded cDNA was prepared from 2ng of RNA using the Nugen Ovation RNAseq System V2. Illumina sequencing libraries were generated from 600 ng of amplified cDNA, after shearing to ~200bp using a Covaris sonicator as described in the protocol (NEBNext Ultra DNA Library Prep Kit for Illumina, #E7370S).

### Western Blots

Three independent replicates of the cortex and SN input samples were obtained. Crude lysate was run on a 4-12% polyacrylamide gel (BioRad) and transferred to PVDF membranes using a semi-dry transblot apparatus (BioRad). Membranes were probed using the following antibodies at the stated concentrations: Chicken GFP (Aves) 1:1000, Mouse Psd95 (Enzo) 1:1000, Mouse Ezrin (Abcam)1:500, LaminB2 (Sigma), 1:500.

### RNA-sequencing

Libraries were sequenced on an Illumina HiSeq 2500, for a total of 175M 50bp reads across the 12 samples. Reads were analyzed as described(45). Briefly, reads were trimmed using Trimmomatic (v0.32), rRNA reads were depleted with bowtie2. Surviving reads were aligned to Ensembl release 75 of the mouse genome using STAR and then counted using HTSeq. Differential expression analysis was performed using the edgeR package. CPMs of all 12 are in Table S1. Raw and analyzed RNA-sequencing data are also available at GEO: GSE74456.

PAP enriched transcripts were determined by the intersect of: transcripts enriched in SN input/Cortex input (FDR < 0.1, Log2(Fold Change) > 1.5) and transcripts enriched in PAP TRAP/SN input (FDR < 0.1, Log2(Fold Change) > 1.5). These transcripts are listed in Table S2.

Soma enriched transcripts were determined by the intersect of: transcripts depleted in SN input/Cortex input (FDR < 0.1, Log2(Fold Change) < 1.5) and transcripts enriched in Cortex TRAP/Cortex Input (FDR < 0.1, Log2(Fold Change) > 1.5). These transcripts are listed in Table S3.

### Gene Ontology Analysis

PAP and Soma lists (see RNA-sequencing and Fig. 3C) were analyzed using the BiNGO plugin(v3.0.3) in Cytoscape(v3.2.1). Mouse Genome Informatics (MGI) IDs and descriptions were obtained from the biomaRt package from Bioconductor in R. Overrepresented biological processes were identified using a hypergeometric test with Benjamini-Hochberg. Lists were compared to a background list of genes detected at CPM > 2 in Cortex input sample.

### Sequence feature analysis

For each gene, the sequence of the longest isoform was obtained using the biomaRt package in R and used in subsequent analyses. We analyzed sequence length, GC content, and individual base content of the 5’UTR, CDS, and 3’UTR, as well as complexity of secondary structure in the 5’UTR and 3’UTR using the Vienna RNA package(46). All comparisons between Soma or PAP lists were made with Wilcoxon rank-sum test with Benjamini-Hochberg correction. To predict RNA structure complexity, we used the ViennaRNA RNAfold program, which calculates the minimum free energy secondary structure of the input RNA sequence. Since longer sequences are more likely by chance to have more complex secondary structures, we normalized this free-energy value by the length of the sequence.

## Author contributions statement

RO developed and KS and RO refined the PAP-TRAP method. KS, AL, and JD designed and conducted bioinformatic analysis. RK, KS and JD conducted ISH, IF, imaging, quantification, and analysis. KS conducted and analyzed puromycylation studies. AD conducted STORM. MV developed image analysis pipeline for quantification. JD and KS conceived of study, designed experiments, and wrote manuscript.

## Acknowledgments

We thank Dr. Joshua Dearborn for help with the neonatal injections, Dr. Min-Yu Sun for guidance in preparing acute slices, Drs Dennis Oakley and James Fitzpatrick for training and assistance with imaging, Dr. Kelly Monk for comments on the manuscript, and the members of the Dougherty lab for advice. This work was supported by the CDI(MD-II-2013-269, and a WUCCI Microgrant), NIH(DA03 8458-01, MH099798-01), WUSTL Interface of Psychology Neuroscience and Genetics Training Grant (5T32 GM081739) and WUSTL Neurosciences Program Training Grant (T32 GM008151). Key technical resources were provided by the Hope Center and the Genome Technology Resource Center at Washington University(supported by NIH grants P30 CA91842 and UL1TR000448), and the Washington University Center for Cellular Imaging(supported by CDI, St. Louis Children’s Hospital, the Foundation for Barnes-Jewish Hospital, and NIH:NS086741).

## Supplemental Figure Legends

**Fig. S1:**
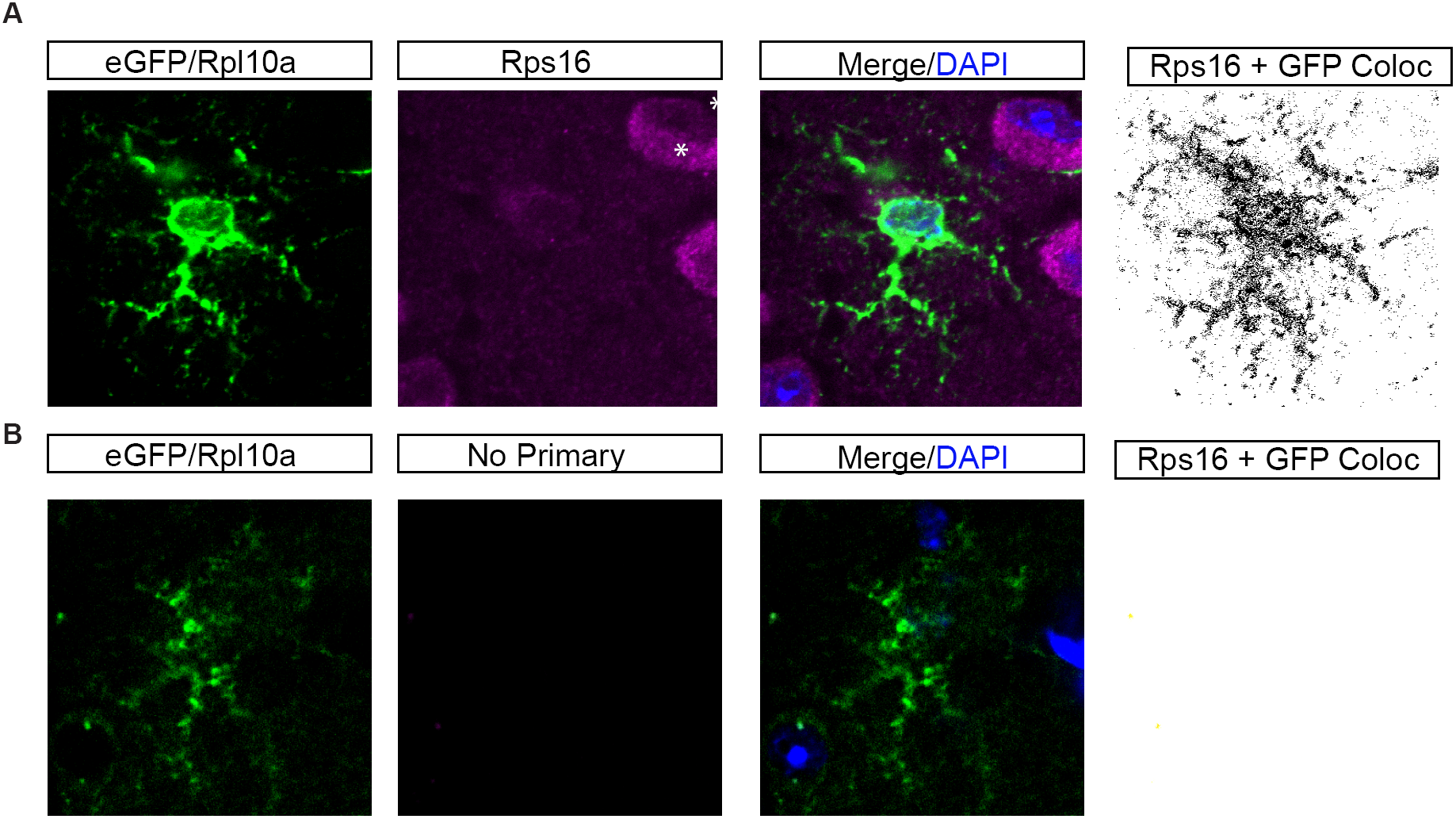
**EGFP/Rpl10a in AldhlLl bacTRAP mouse cortex extends beyond Gfap+ processes** (A) Examination of localization of Rps16 by confocal reveals detectable Rps16(violet/white) in fine processes. Note Rps16 is also found robustly in adjacent neurons, as expected (*). Therefore, presence of GFP signal was used to mask all non-astrocyte pixels, and a binary image created showing GFP positive pixels that also contained Rps16 signal. This approach allows for identification of Rps16 labeling specifically in the astrocyte even in the presence of Rps16 labeling in adjacent cells. (B) A tissue section processed in parallel, but omitting Rps16 primary antibody, demonstrates absence of signal in the astrocyte.

**Fig. S2:**
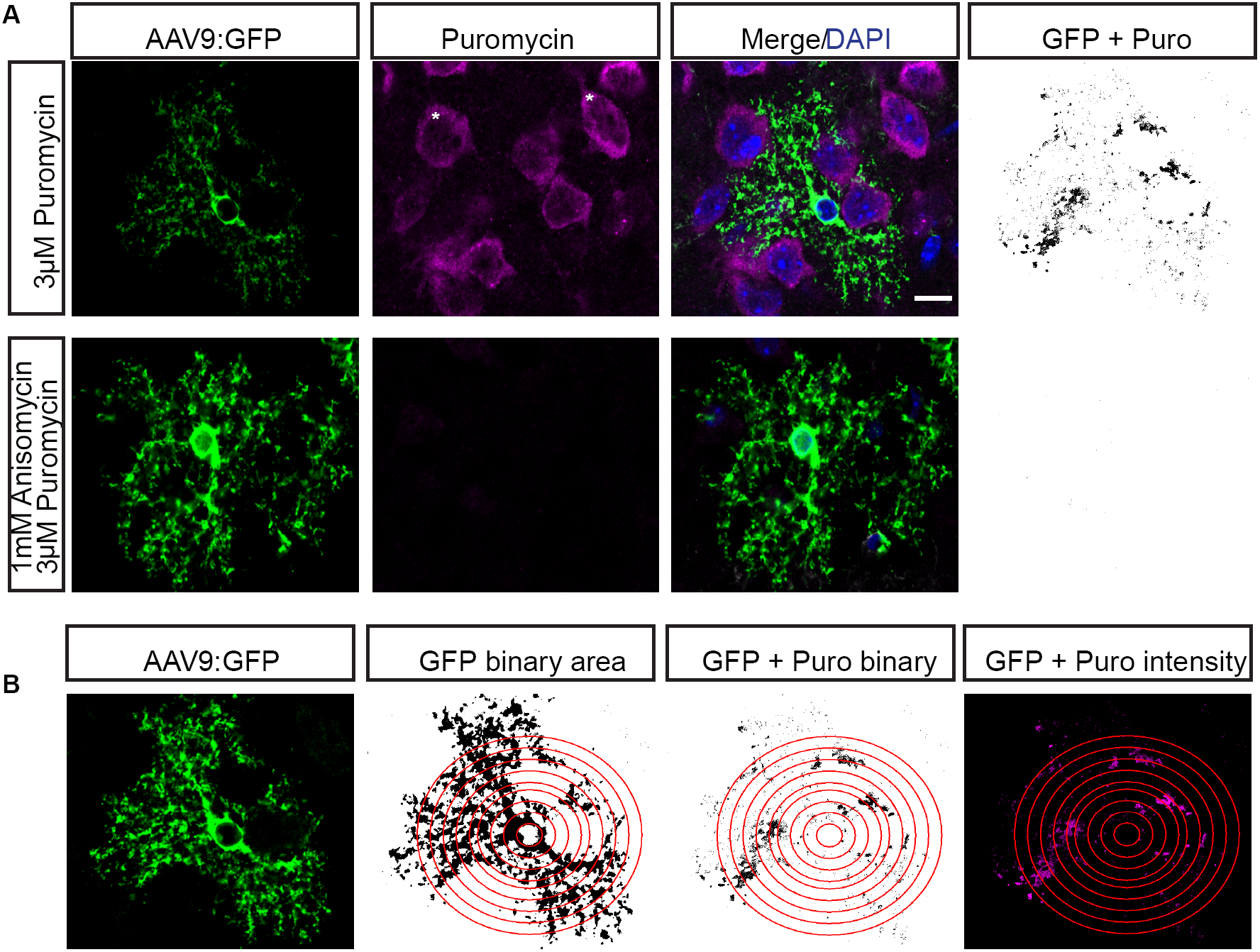
**Puromycylation detection and quantification method.** (A) Example of masking for quantification of peripheral puromyclation (Confocal IF). Note, as expected, robust translation also occurs in adjacent neurons(*).Therefore, presence of GFP signal was used to mask all non-astrocyte pixels, and a binary image created showing GFP positive pixels that also contained Puro-IF signal (GFP+Puro) for all downstream quantification. (B) Illustration of method for quantification of relative amount of translation at increasing distance from the astrocyte nucleus. A nucleus of a GFP+ astrocyte is selected and thresholded (middle left). A series of Sholl-like concentric rings are demarked for quantification (red rings). All Puro-IF outside of GFP+ area is masked, and Puro-IF positive area within each ring is measured (middle right). Intensity of Puro-IF within each GFP+ pixel is also measured (right panel).

**Fig. S3:**
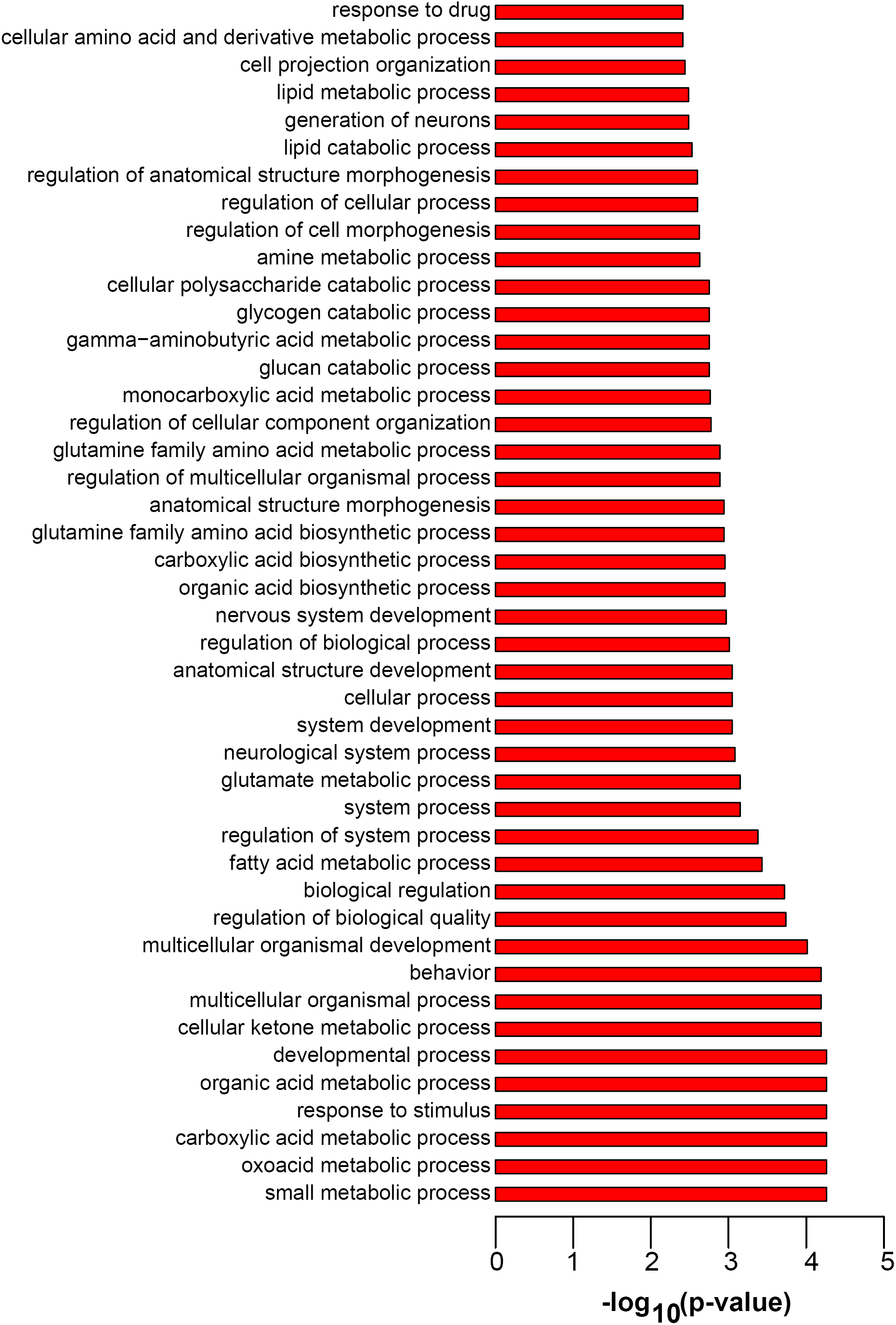
**Pathway analysis using Gene Ontologies for PAP enriched transcripts.** All enriched GO terms for PAP-enriched transcripts, p < 0.005, hypergeometric test with B-H correction.

**Fig. S4:**
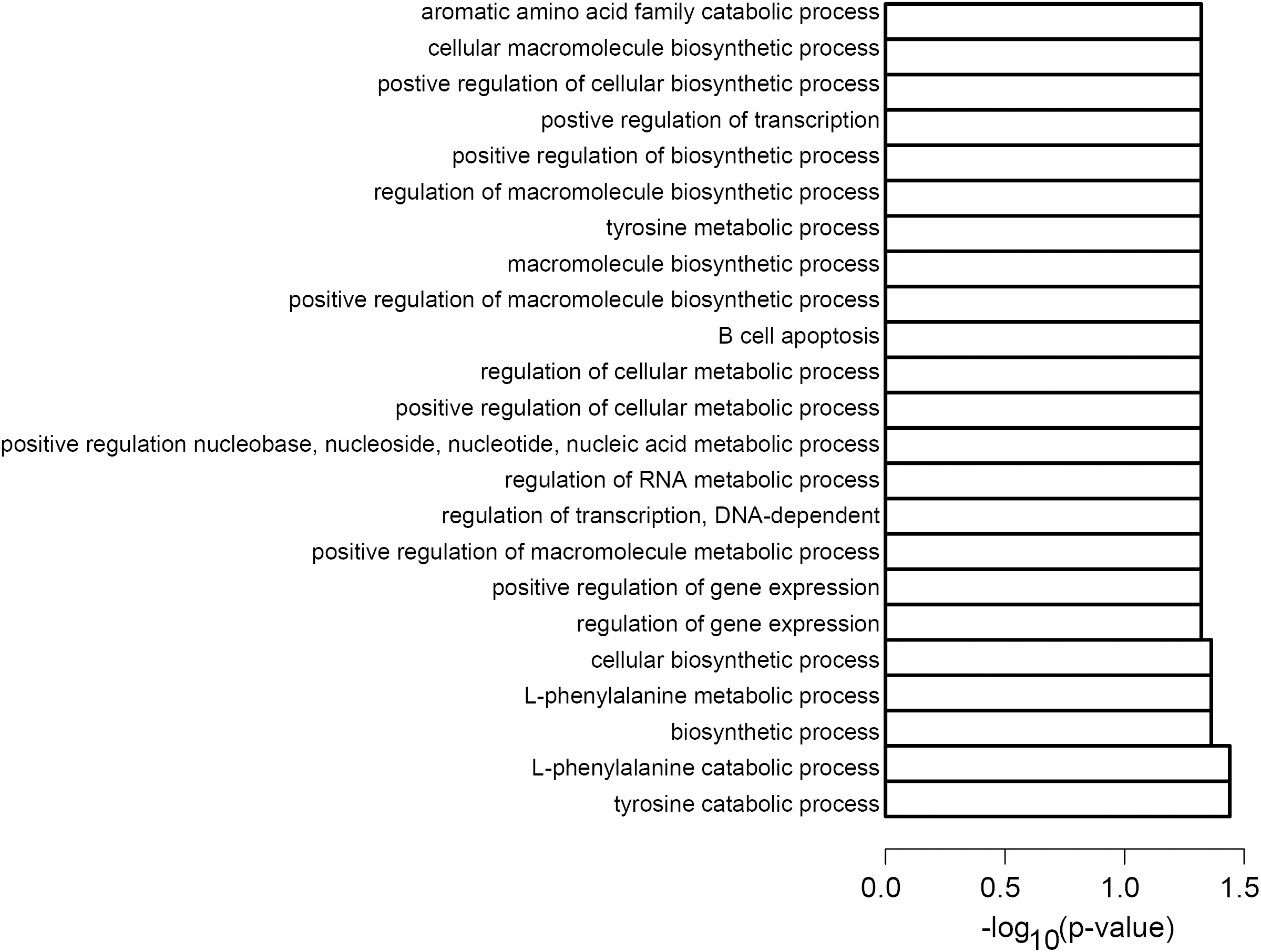
**Pathway analysis using Gene Ontologies Soma enriched transcripts.** All enriched GO terms for Soma-enriched transcripts, p < 0.05, hypergeometric test with B-H correction.

**Fig. S5:**
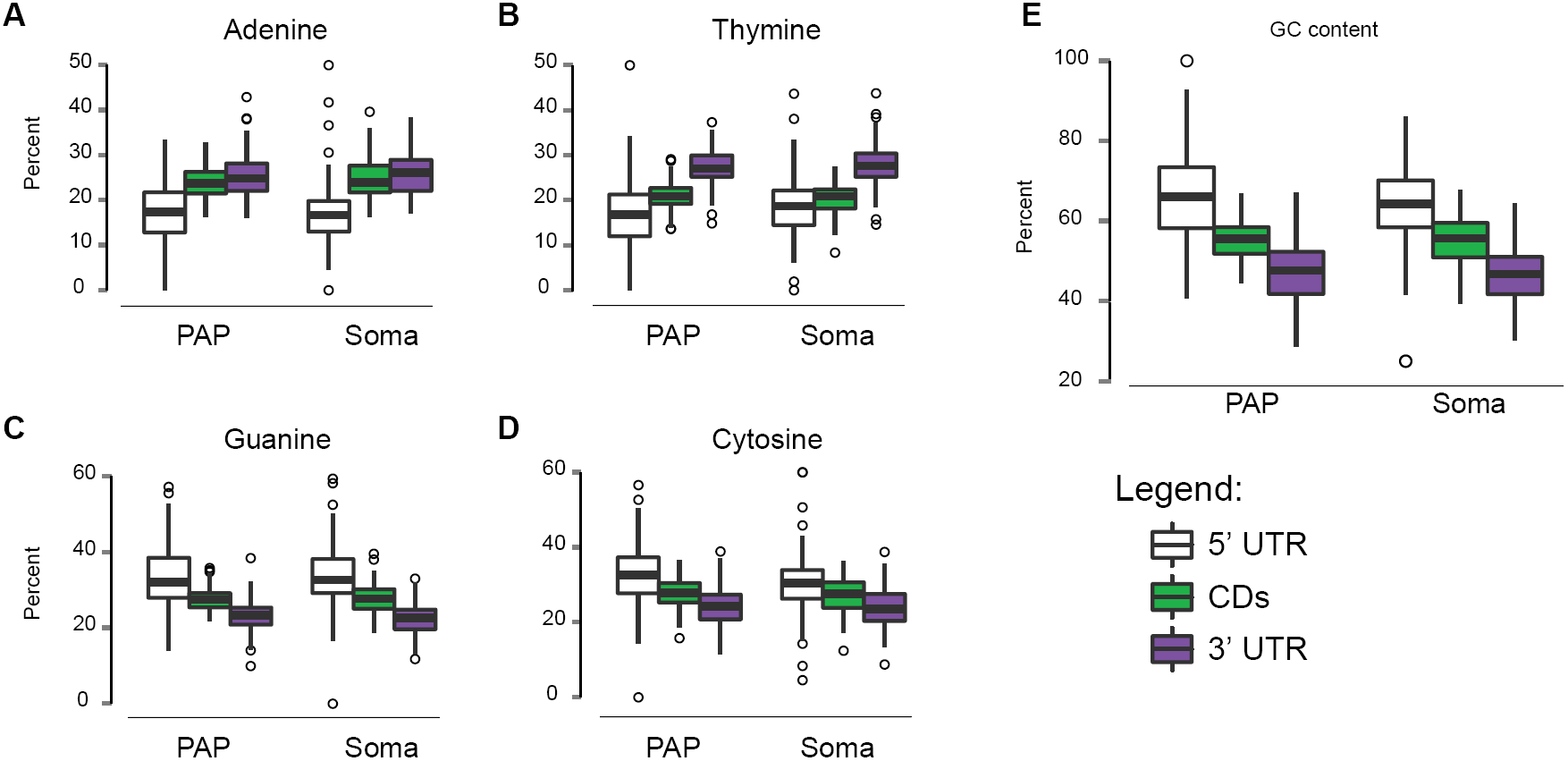
**Nucleotide composition does not differ between PAP and Soma-enriched transcripts. (**A-D) Percent composition of each nucleotide for 5’ UTR (white), coding sequence (CDs, green), and 3’UTR (purple) in PAP and Soma lists. p > 0.05, Wilcoxon test with B-H (E) GC percentage. p > 0.05, Wilcoxon test with B-H correction.

**Fig. S6.**
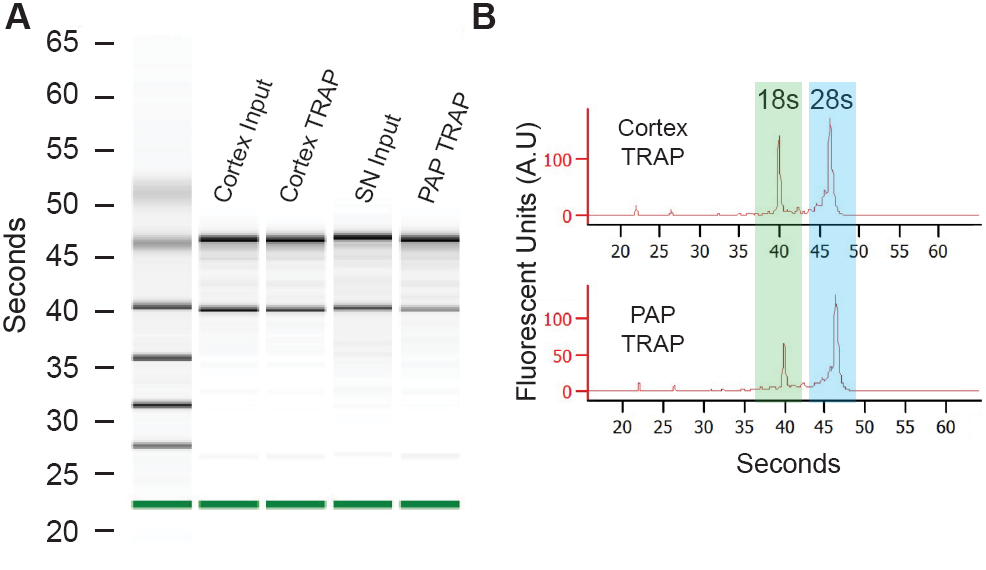
**Bioanalyzer confirms RNA quality from Cortex TRAP and PAP TRAP samples.** (A) PicoChip electrophoresis for representative samples, marker is highlighted in green. (B) Fluorescent trace visualization of PicoChip electrophoresis for Cortex TRAP and PAP TRAP. Ribosomal RNAs from both subunits are captured.

**Fig. S7.**
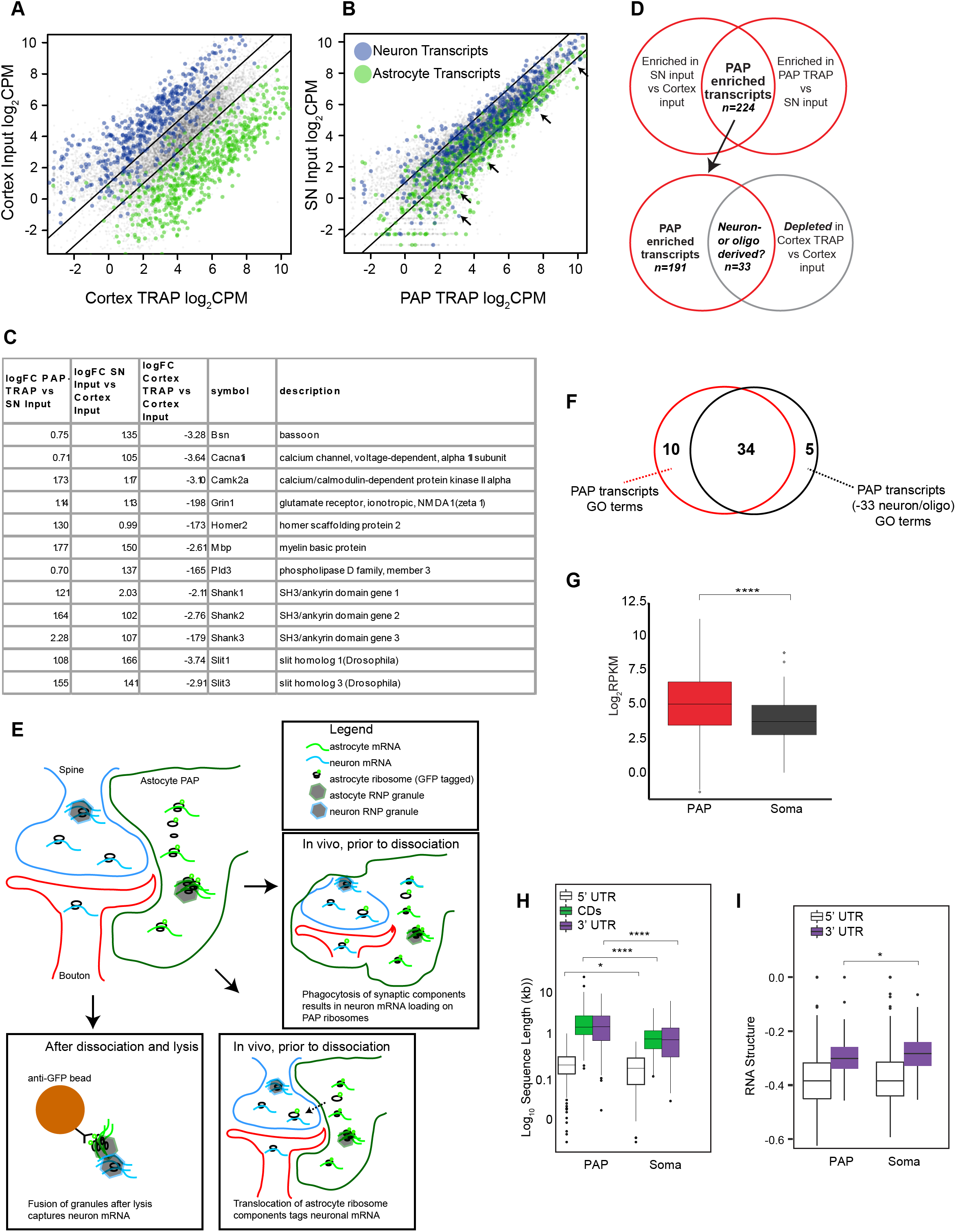
**A small number of neuronal transcripts are bound to peripheral astrocyte ribosomes: in vitro or in vivo mechanism?** (A) Scatterplot of Cortex TRAP compared to Cortex Input CPM for all transcripts (Log2). We detect robust enrichment for sets of ~ 200 previously detected(1) astrocyte-enriched transcripts (green dots), and substantial, though not complete, depletion of ~200 previously detected(1) neuron-enriched transcripts (blue dots) in the current cortex TRAP experiment. This degree of enrichment and depletion is typical for TRAP(2, 3) (B) While PAP-TRAP still enriches for most astrocyte transcripts relative to SN samples, scatterplot of PAP-TRAP compared to SN input shows an unexpected relative blunting of both enrichment and depletion relative to Cortex TRAP. Note, significantly enriched transcripts are still detectable, as evidenced in Table S2. Surprisingly, a handful of ‘neuronal’ transcripts now show relative enrichment following capture of astrocyte ribosomes (arrows) from the SN fraction (PAP-TRAP). (C) These include examples of many transcripts that are strong candidates to be locally translated in neurons (*Camk2a, Shanks*), or are known to be locally translated in oligodendrocytes (e.g., *Mbp*)(4). (D) Roughly 15% of PAP-Enriched transcripts from Table S2, which show clear enrichment by PAP-TRAP, are depleted when examining the Cortex TRAP vs Input comparison, suggesting they are not present on most astrocyte ribosomes in the cell body, as illustrated by Venn Diagram. (E) Illustrations of some of the possible alternate models for these unexpected results. Left panel: interactions in solution between RNA granules originally from neurons with captured granules from astrocytes *in vitro,* mediated by the disordered nature of domains in many RBPs(5). Right panels: low level exchange of mRNA or ribosomal components between adjacent processes, or perhaps following astrocyte phagocytosis of synapses(6), *in vivo.* Regardless, if needed, this 15% can be removed from the PAP-Enriched list computationally (provided as Table S4), and all the major conclusions regarding the remaining PAP-Enriched genes are still supported. (F) 77% of the original PAP GO terms survive at B-H corrected p<0.005. (G) Quantification of expression of PAP-enriched and Soma-enriched transcripts indicates Soma transcripts have lower median expression in cortical astrocytes(Wilcoxon test, B-H corrected, *p<0.0001). (H) Quantification of length of PAP and Soma transcripts indicates PAP-enriched transcripts are longer (Wilcoxon test, B-H corrected, ****p<0.0001, *p<0.05). (I) RNA structure-score(minimum free energy of most stable predicted structure, normalized to length), indicates PAP-enriched transcripts have more stable 3’UTR secondary structures(Wilcoxon test, B-H corrected, *p<0.05, lower values are more stable)

**Table S1. CPM** Counts per million for all biological replicates. Nomenclature: Cortex input = cortex homogenate transcripts, Cortex TRAP = transcripts on GFP+ ribosomes in Aldh1l1:eGFP/Rpl10a mouse cortex, SN input = transcripts in SN fraction, PAP TRAP = transcripts on GFP+ ribosomes in Aldh1l1:eGFP/Rpl10a SN fraction.

**Table S2. PAP enriched transcripts** PAP-enriched transcripts, as defined in Figure 4, with respective sample enrichment and FDR significance. Log fold changes (base 2) and FDR for three comparisons are included in the table. Abbreviations as in Table S1.

**Table S3. Soma enriched transcripts** Soma-enriched transcripts, as defined in Figure 4, with respective sample enrichment and FDR significances, as in Table S2.

**Table S4. PAP enriched transcripts, with putative neuron or oligodendrocyte derived transcripts removed.**

PAP-enriched transcripts, as defined in S7, with 33 transcripts removed that were significantly depleted (FDR<.1, logFC<-1) in Cortex TRAP vs Cortex input comparison.

